# Dry Swab Method of sample collection for SARS-CoV2 testing can be used for culturing virus

**DOI:** 10.1101/2021.03.23.436593

**Authors:** Sushma Ram, M. Ghalib Enayathullah, Yash Parekh, Karthik Bharadwaj Tallapaka, Rakesh K Mishra, Kiran Kumar Bokara

## Abstract

**Back ground:** Earlier studies suggested the use of dry swab method for SARS-CoV-2 detection as it does not need VTM and subsequent RNA extraction step making the process cheaper, safer and faster. In this study we explore whether the virus in the dry swab is viable and can be cultured and propagated.

**Method:** Swabs were spiked with SARS-CoV-2 and stored in three different conditions: a) as dry swab (S_D_, eluted in 1 mL DMEM), b) in 1 mL of Viral Transport Medium (S_VTM_), and c) in 1 mL of Tris-EDTA buffer (S_TE_). The sample groups were stored either at room temperature (RT, 25°C±1°C) or at 4°C for 1, 4, 8, 12, 24, 48 and 72 hours before being used as viral inoculums for the propagation studies in Vero cells.

**Results:** The RT-qPCR data suggests that S_D_ incubated both at RT and 4°C harbors viral particles that are viable and culturable at par with S_VTM_ and S_TE_.

**Conclusion:** The dry swab method, in addition to its advantages in detection of the virus, also renders viable viral particles that can be cultured and propagated.

## 1. Introduction

Corona Virus Disease (COVID-19), originating in Wuhan of China has wreaked havoc all over the world following its rapid advancement into a pandemic. Currently, the most common method of Severe Acute Respiratory Syndrome Corona Virus 2 (SARS-CoV-2) detection in clinical settings is an RT-qPCR assay following RNA isolation from patients’ respiratory samples. The present-day necessity for more economic, effective and rapid detection methods has resulted in usage of various modifications of the technique like elimination of transport medium and RNA isolation step^1,2,3,4^. It has been reported that there is no significant difference between C_t_ values of swab samples incubated in dry conditions and those incubated in liquid media^5^.

While these methods diminish the testing time and expenses, the viability of the virus in samples procured by dry swab method, for culturing, has not been established. In the present study, we demonstrate that viable viral isolates can be obtained and propagated from the dry swab sample collection method.

## 2. Materials and Methods

### 2.1 SARS CoV2 Viral Culture

The established culture of SARS-CoV-2 virus in our institute (Indian/a3i clade/2020 isolate) was used for the present study.

### 2.2 Sample preparation and Processing

The viral stock of a known titer was diluted at 1:2 ratio in DMEM culture medium (without FBS) and 40 microliters (~10^5^ viral particles) of the viral inoculum was used to spike the swabs from the experimental groups S_D_, S_VTM_ and S_TE_. The sample groups were incubated at room temperature (25°C±1°C) and at 4°C for 1, 4, 8, 12, 24, 48 and 72 hours. After the incubation, all samples were stored at −80°C until further use.

### 2.3 SARS CoV2 Viral Infection

Vero cells seeded in 96-well plates were cultured at 37°C, 5% CO_2_ in Dulbecco Minimum Essential Medium (DMEM) (Gibco) supplemented with 10% (v/v) Fetal Bovine Serum (FBS) (Gibco), 3.7 g/L sodium bicarbonate. At 90-95% confluency, the cells were infected with virus from S_D_ (eluted in 1 mL DMEM), S_VTM_ and S_TE_ diluted at 1:5 ratio in DMEM culture media (without FBS) for 3 hours. Later, the virus containing medium was aspirated and was replaced with fresh DMEM with 10% FBS. Samples of uninfected Vero cells and those infected with viral stock diluted 1:10 (~MOI 0.1) were included as cell control and infection control, respectively.

### 2.4 RNA Isolation

RNA was extracted from 200 μL aliquots of culture supernatants using the MagMAX™ Viral/Pathogen Extraction Kit (Applied Biosystems, Thermofisher). Extraction of viral RNA was carried out according to the manufacturer’s instructions. The viral supernatants from the test groups were added into the deep well plate (KingFisher™ Thermo Scientific) along with a lysis buffer containing the following components - MagMAX™ Viral/Pathogen Binding Solution; MVP-II Binding Beads; MagMAX™ Viral /Pathogen Proteinase-K of 260 μL; 10 μL; 5 μL respectively for 200μL of sample. RNA extraction was performed using KingFisher Flex (version 1.01, Thermo Scientific) by following manufactures instructions. The eluted RNA was stored at −80 μC until further use.

### 2.5 RT-qPCR for detection of SARS CoV2

ICMR approved Meril Covid-19 one step RT PCR kit was used to detect the ORF 1ab (FAM labeled) and nucleoprotein N (HEX labeled) genes of SARS-CoV2 in the isolated RNA samples. Reaction was set up according to manufacturer’s protocol: Reverse Transcription-50 ^0^C, 15 minutes; cDNA Initial Denaturation 95 ^0^C, 3 minutes; 45 cycles of Denaturation at 95 ^0^C, 15 seconds and Annealing, Extension and Fluorescence measurement at 55 ^0^C, 40 seconds; Cooling-25 ^0^C, 10 seconds. The program was set up using QuantStudio-5 machine (Thermo fisher). The C_t_ values of N gene and ORF1ab (genes specific to SARS-CoV-2) were considered to plot the graphs.

### 2.6 Statistical analysis

The RT-qPCR data are mean of at least three independent experiments. Data is represented as mean ± S_D_ using GraphPad Prism 8 (Ver 8.4.2 GraphPad Software, LLC.).

#### Ethics approval statement

The Anti-SARS CoV2 study was approved from Institutional Bio-safety Committee of CSIR-Center for Cellular and Molecular Biology, Hyderabad, India

## 3. Results and Conclusions

The present study uses known SARS-CoV-2 inoculum to spike experimental swabs in order to mimic the conditions of sample collection from patients. Our results showed that the viral elutes from swab samples of S_D_, S_VTM_ and S_TE_ experimental groups exhibited similar viability both at 25°C and at 4°C as evidenced by the C_t_ values of N gene and ORF1 ab gene obtained from the RT-qPCR (Figure 1 & 2).

**Figure 1:**
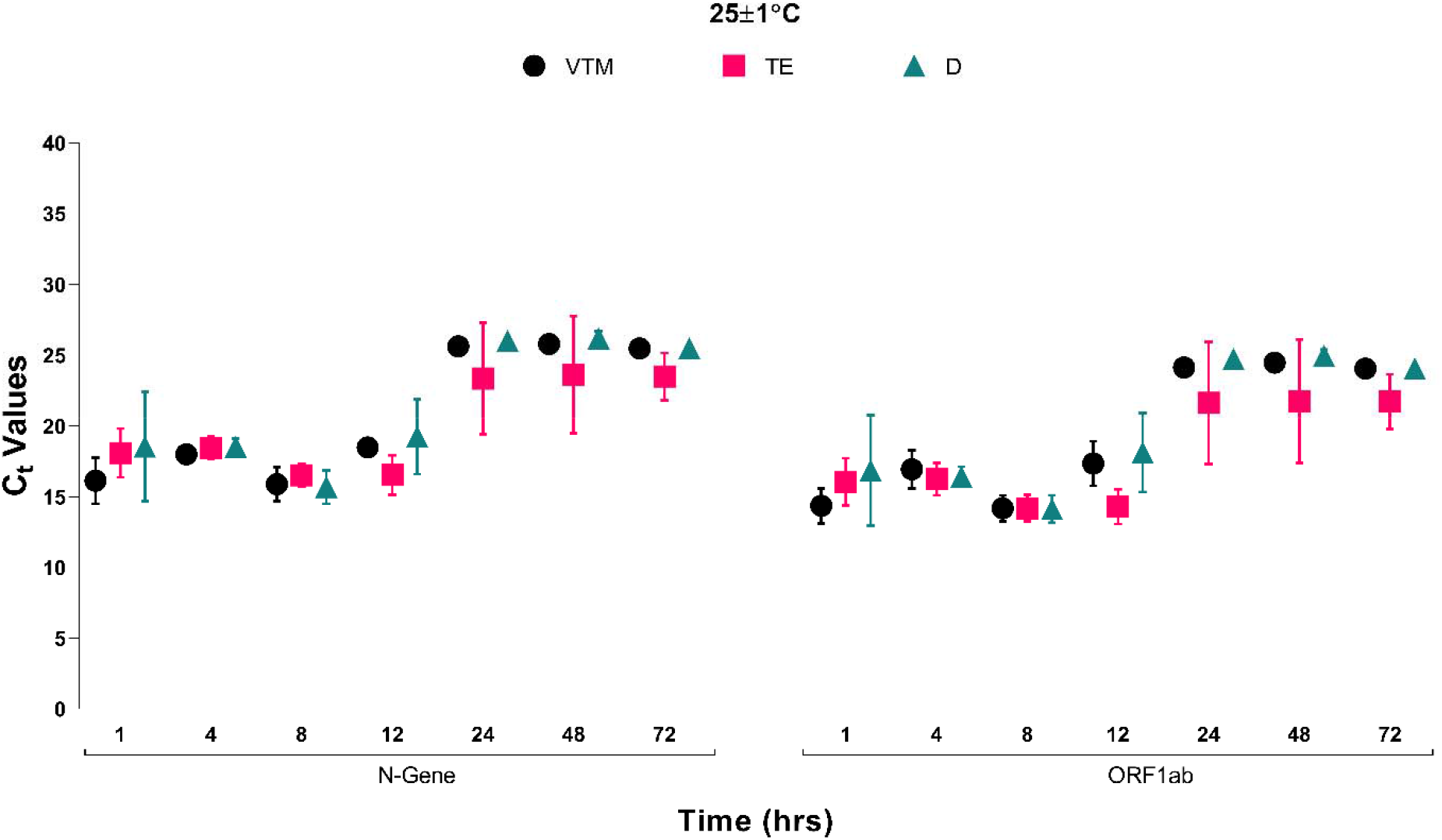
The plot represents the C_t_ values of N gene and ORF 1ab genes of SARS CoV2 virus obtained by RT-qPCR from culture supernatants of S_D_, S_VTM_ and S_TE_ infected samples incubated at 25°C for 1, 4, 8, 12, 24, 48 and 72 hours

**Figure 2:**
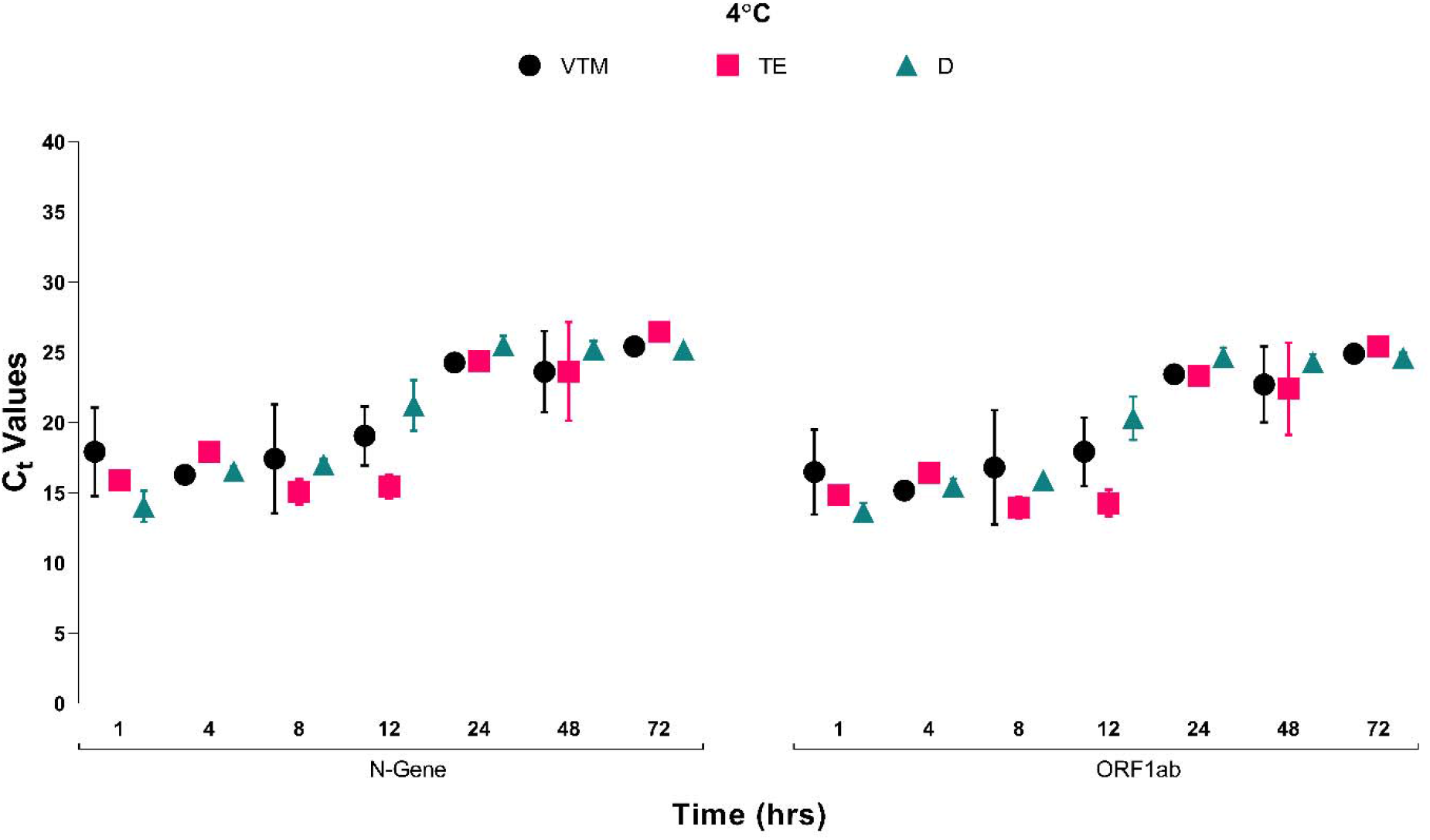
The plot represents the C_t_ values of N gene and ORF 1ab genes of SARS CoV2 virus obtained by RT-qPCR from culture supernatants of S_D_, S_VTM_ and S_TE_ infected samples incubated at 4°C for 1, 4, 8, 12, 24, 48 and 72 hours.

The C_t_ values ranged between 16 and 19 in S_D_, S_VTM_ and S_TE_ up to 12 hours either at 25°C or 4°C. However, slight increase in the C_t_ values were observed over 24 hours onwards until 72 hours and the C_t_ values ranged between 23-26. This indicates that with increase in time, the viability of the viral particles is compromised and hence early retrieval of the viral particles either from S_D_ or from S_VTM_ and S_TE_ is recommended. Our results also suggest that the retrieval of viable viral particles from S_TE_ is slightly better (~ −2 C_t_) than that from S_D_ or from S_VTM_.

While the dry swab approach for sample collection is beneficial economically and for hastening diagnosis, it also helps avoid liquid media that may interfere with RT-qPCR detection^3^. The utilization of Dry swab may simplify sample collection, eliminate the need for liquid transport media and facilitate the retrieval of viable viral particles for culturing and subsequent use in research.

## Acknowledgements

We thank Mr Uday Kiran, Mr C.G Gokulan and Mr Santosh Kumar Kuncha for providing technical support for dry swab processing.

